# Genomic Characterisation of the Irish Traditional BóRiabhach Cattle Breed

**DOI:** 10.64898/2026.11.21.768272

**Authors:** Darren McLoughlin, Gillian P. McHugo, Paul Flynn, Patrick Sullivan, Sam S. Browett, David E. MacHugh, James A. Ward

**Affiliations:** UCD School of Agriculture and Food Science, University College Dublin, Belfield, Dublin D04 V1W8, Ireland; BóRiabach Cattle Society, Gurteen, Carrigallen, Leitrim H12 F895, Ireland; Irish Rare Breeds Society, Gurteen, Carrigallen, Leitrim H12 F895, Ireland; Weatherbys Scientific, Naas, Kildare, W91 VX86, Ireland; Department of Land Sciences, South East Technological University (SETU), Waterford, Waterford, X91 K0EK, Ireland; UCD Conway Institute of Biomolecular and Biomedical Research, University College Dublin, Belfield, Dublin D04 V1W8, Ireland; UCD One Health Centre, University College Dublin, Belfield, Dublin D04 V1W8, Ireland; Chair of Animal Systems Genomics, Faculty of Veterinary Medicine, Ludwig-Maximilians-Universität, 80539 Munich, Germany

**Keywords:** cattle, conservation genomics, endangered breed, inbreeding, genetic diversity, population genomics, selection signature, single nucleotide polymorphism

## Abstract

The BóRiabhach is a highly endangered traditional landrace cattle breed in Ireland with a culturally significant history. The Irish Gaelic word “*riabhach*” means “streaked”, “striped”, or “brindled”, and a distinctive brindle coat colour pattern is characteristic of the BóRiabhach breed. The breed had declined to about 20 animals by 2016 and has no systematic pedigree. Despite conservation efforts that have increased the population to an estimated 70 animals, the breed remains one of Europe’s most critically endangered livestock genetic resources. We present the first genomic characterisation of the BóRiabhach breed, using Illumina BovineSNP50 genotypes of 84 BóRiabhach animals and 688 cattle from 28 European comparator breeds. Using model-based clustering, we identified a BóRiabhach-specific ancestry component. Only 28 animals carried more than 50% of this component, which we term “*high*-BRBH”; the remaining 56 showed extensive introgression, chiefly from Limousin and Holstein, which explains the breed’s high observed heterozygosity. Phylogenetic analysis placed the *high*-BRBH cohort with British Shorthorn and Lincoln Red within a British-Irish clade, distinct from Kerry and Dexter, and recovered a Limousin-to-BóRiabhach migration edge. This *high*-BRBH core group exhibited the lowest effective population size (*N*_e_) of any adequately sampled breed and consists of closely related animals: 34% of pairs within the core were second-degree relatives or closer. Conservation of the BóRiabhach must therefore balance breed distinctiveness with genetic diversity. We recommend genotyping all extant animals, recording genomic ancestry at registration, managing mating based on genomic relatedness as well as ancestry, and cryopreserving germplasm from the *high*-BRBH core population.

## Introduction

Cattle have been domesticated at least twice; first in the Fertile Crescent, taurine cattle (humpless *Bos taurus*) were domesticated over 10,000 years ago from wild aurochs (*Bos primigenius primigenius*), and second, approximately 2,000 years later, zebu cattle (humped *Bos indicus*) were domesticated in South Asia from a different aurochs subspecies (*B. p. namadicus*) (Larson & Fuller 2014; Larson *et al*. 2014; McHugo *et al*. 2019b; Pitt *et al*. 2019). Beginning in the mid-20^th^ century, farmers favoured highly productive dairy and beef cattle breeds, particularly in the industrialised economies of Europe, North America, and Australasia. This process has accelerated during the last several decades and has led to the extinction and increased vulnerability of more than 200 traditional landrace or heritage cattle breeds across Europe (Scherf & Pilling 2015). The rapid loss of cattle genetic resources has become all the more pressing as climate change accelerates and agroecological environments are transformed, particularly in northern Europe (Hoffmann 2010). Therefore, traditional landrace breeds represent important reservoirs of genetic variation for cattle breeding programmes to address future agroecological and socioeconomic challenges, including those that will leverage genomic technologies such as genomic selection and genome editing (Georges *et al*. 2019; Rochus *et al*. 2025).

The BóRiabhach is a highly endangered traditional landrace cattle breed in Ireland with a culturally significant history. The Irish Gaelic word “*riabhach*” means “streaked”, “striped”, or “brindled”, and its spelling has remained unchanged since the Old Irish of the Early Medieval Period (Ó Dónaill 1977). A distinctive brindle coat colour pattern is characteristic of the BóRiabhach breed (**Figure 1A, 1B**). The BóRiabhach breed experienced a dramatic population decline in the 20^th^ century, and by 2016, only about 20 BóRiabhach cattle were estimated to survive. Conservation efforts by individual breeders and the formation of the BóRiabhach Cattle Society in 2019 have helped increase the population to an estimated 70 animals; however, the breed remains one of Europe’s most critically endangered livestock genetic resources (BóRiabhach Cattle Society 2026; Food and Agriculture Organization of the United Nations 2026). The conservation of endangered breeds, such as the BóRiabhach, is supported by the FAO Global Plan of Action for Animal Genetic Resources and, within the European Union, by Regulation (EU) 2016/1012 (Food and Agriculture Organization 2007; European Parliament and Council of the European Union 2016).

**Figure 1:**
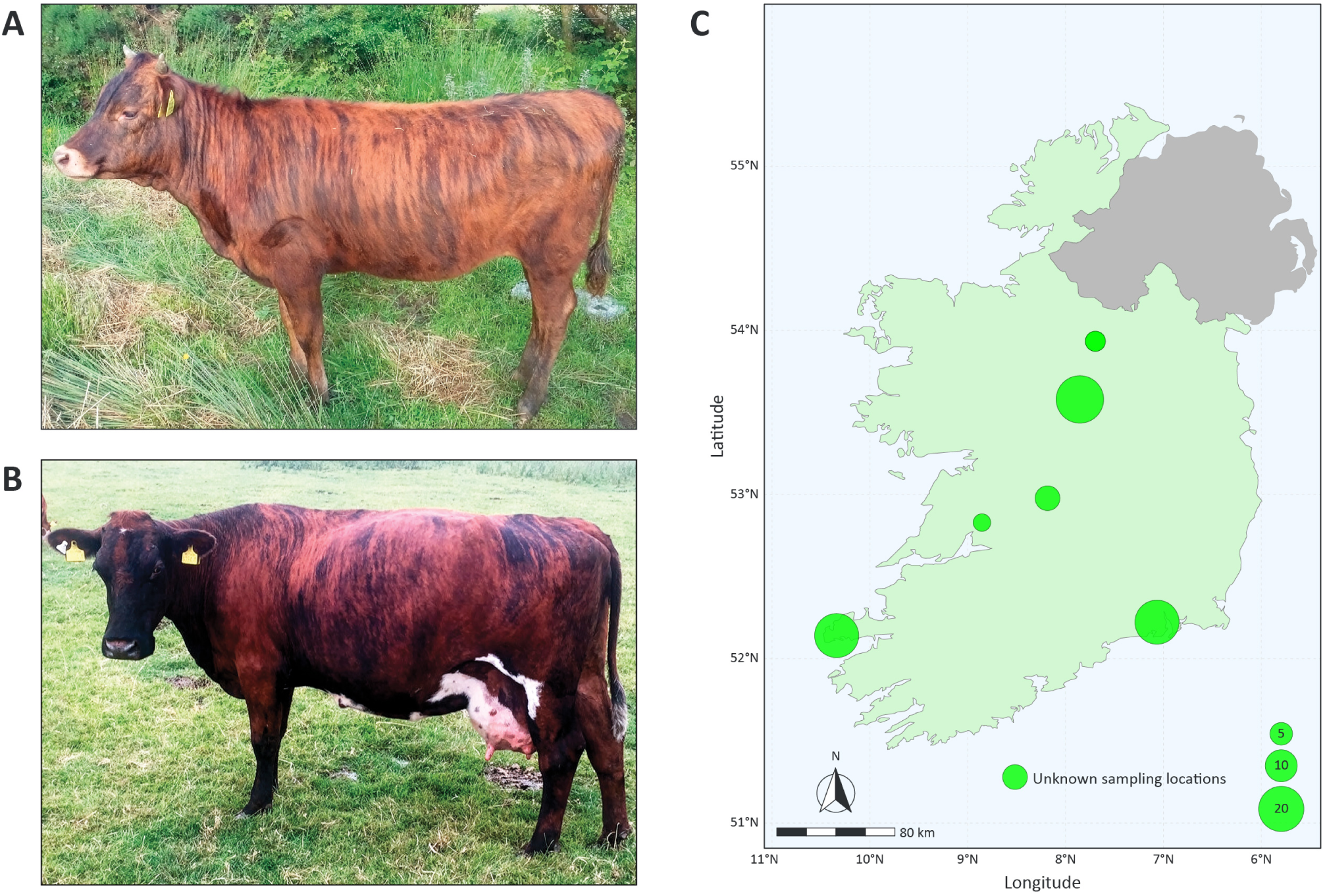
Representative BóRiabhach female animals. (**A**) A female BóRiabhach calf showing a developing brindle coat colour pattern. (**B**) An adult BóRiabhach cow showing the fully developed brindle coat colour pattern. ©Darren McLoughlin (2019, 2021). (**C**) Locations of BóRiabhach cattle herd sampling in the Republic of Ireland for the 84 animals used in the present study. The area of each circle corresponds to the number of animals sampled from each herd (as indicated in the reference key). Six of the 84 cattle had unknown sampling origins, which are also indicated off the map.

Livestock conservation genetics has been revolutionised by rapid, cost-effective technologies enabling high-throughput, genome-wide genotyping of single-nucleotide polymorphisms (SNPs). These methods have enabled comprehensive, increasingly fine-grained assessments of genetic diversity, population structure, inbreeding levels, genome-wide signatures of selection, and demographic history in threatened livestock populations (Browett *et al*. 2018; Ajmone-Marsan *et al*. 2023). For the present study, to provide an evidence base for genetic conservation, we conducted high-resolution comparative population genomics analyses of BóRiabhach cattle and a range of European cattle breeds, informed by previous work on the conservation genomics of traditional Irish livestock (Browett *et al*. 2018; McHugo *et al*. 2019a) Consequently, this study focused on reconstructing phylogenetic networks, assessing genetic structure and inbreeding, modelling historical population sizes, and investigating artificial and natural selection across the BóRiabhach genome.

## Materials and Methods

### Samples and Genotyping

We obtained genomic data from the BóRiabhach Cattle Society, collected at owners’ request for routine genetic improvement and population monitoring. We also collated published genomic data for all comparator breeds. (Browett *et al*. 2018). For the present study, an Illumina^®^ BovineSNP50 BeadChip (Matukumalli *et al*. 2009) dataset was used, comprising 84 individual animals from the BóRiabhach (BRBH) cattle population that were genotyped by Weatherbys Scientific (Co. Kildare, Ireland) using DNA samples purified from hair follicles and nasal swabs. **Figure 1C** shows the sampled BRBH animals and their herd-of-origin locations. Comparator BovineSNP50 datasets comprising a total of 710 individual *B. taurus* animals and representing a total of 29 different breeds from Ireland, Britain, Europe, and West Africa (taurine N’Dama from Guinea – NDAG) were obtained from the *WIDDE* resource (Sempéré *et al*. 2015) and the breed/population sample datasets detailed by Browett *et al*. (2018). **Supplementary Table 1** presents the sample sizes and additional biogeographical and production information for all 30 breeds used in this study.

### Genome assembly SNP updating, quality control, and filtering

The complete BovineSNP50 dataset for all 30 breeds (794 animals: 772 European cattle and 22 NDAG) comprised 54,057 SNPs before SNP quality control (QC) and filtering. SNP genomic locations were updated from the UMD3.1 bovine genome assembly to the current ARS-UCD1.2 assembly (Schnabel 2018; Rosen *et al*. 2020) using PLINK (v.1.90 beta 7.11) (Chang *et al*. 2015). We performed SNP QC using PLINK, leaving 42,124 SNPs. Because the analyses have different requirements, we applied SNP filters per analysis; the filters, SNP counts, and animal counts for each are listed in **Supplementary Table 2**. In brief, we removed autosomal SNPs with call rate < 95% throughout; we applied a MAF ≥ 0.05 filter and LD pruning (*r*² < 0.1, 50-SNP window, 10-SNP step) for ADMIXTURE, PCA, and relatedness; we applied no MAF filter or pruning for ROH detection.

### Genetic structure and multivariate principal component analysis (PCA)

We ran ADMIXTURE v1.3.0 (Alexander *et al*. 2009) on 772 animals from 29 breeds (the NDAG were excluded) using 13,653 LD-pruned SNPs, for *K* = 2 to 14 with 10-fold cross-validation. At each *K*, we considered a component BóRiabhach-specific if the BRBH animals had the highest mean of any breed, no other breed averaged > 0.10, no non-BRBH animal exceeded 0.50, and at least five BRBH animals exceeded 0.50. The lowest *K* value meeting these criteria was *K* = 9. We assessed stability at higher *K* by matching components across *K* by correlation. We classified animals with > 0.50 of this component as *high*-BRBH (*n* = 28) and the remainder as *low*-BRBH (*n* = 56). We used the *high*-BRBH group for TreeMix and *N*_e_ estimation, and we analysed both groups for *F*_ROH_ and relatedness. PCA was performed in PLINK on the same animals and SNPs. Figures were produced with ggplot2 v3.5.0 (Wickham 2016).

### Construction of phylogenetic trees with ancestry graphs

To infer the topology of population splits and gene flow among the 30 sampled taurine cattle breeds, we used the TreeMix (v.1.13) program (Pickrell & Pritchard 2012) with autosomal SNP data from 738 animals. We ran TreeMix with 25,395 autosomal SNPs and migration edges ranging from *m* = 0 to *m* = 8, with 10 independent replicates per *m* value, each initialised with a different random seed. Each run used a SNP block size of *k* = 500 to account for linkage disequilibrium (LD) in jackknife variance estimation, with the West African taurine NDAG specified as the root population. The optimal number of migration edges was evaluated using R (v.4.3.2) (R Core Team 2025).

### Modelling recent and historical effective population size (*N*_e_)

We modelled recent and historical *N*_e_ trends using genome-wide autosomal SNP linkage disequilibrium (LD) data for *high*-BRBH and all European breeds, using the SNeP (v.1.11) software tool as described by Barbato *et al*. (2015) and following the procedures detailed by Browett *et al*. (2018). We also used the SNeP program option for unphased SNP data for these analyses.

### Estimation of genomic inbreeding using a runs of homozygosity (ROH)-based measure (*F*_ROH_)

Genomic inbreeding was evaluated as genome-wide autozygosity, estimated from the genome-wide autosomal SNP data using runs of homozygosity (ROH) and the *F*_ROH_ statistic (McQuillan *et al*. 2008). We calculated the *F*_ROH_ statistic as the ratio of the total length of defined ROH (*L*_ROH_) to the total length of the autosomal genome covered by the SNPs for each animal.

The PLINK software toolset was used to define ROH using a sliding-window approach and previously described methods suitable for BovineSNP50 datasets (Purfield *et al*. 2012), which were modified from those provided by Browett *et al*. (2018). The ROH parameters were: 50-SNP sliding window with ≤ 1 heterozygous and ≤ 5 missing calls; minimum 50 SNPs and 1,000 kb; ≥ 1 SNP per 100 kb; and maximum gap 1,000 kb (Purfield *et al*. 2012). The SNP QC steps are described in **Supplementary Table 2**. *F*_ROH_ was the summed ROH length divided by the 2,458 Mb of autosomes covered by SNPs. With a mean inter-SNP distance of 95 kb, the shortest detectable ROH was ≈ 4.7 Mb, so *F*_ROH_ reflects inbreeding over roughly the last 10 generations. ROH were classed as < 8, 8–16, and > 16 Mb, corresponding to common ancestors more than six, three to six, and fewer than three generations ago (for further background on these estimates see: Curik *et al*. 2014).

### Analysis of genomic relatedness in BóRiabhach breed using an identity-by-state (IBS) matrix

We calculated pairwise identity-by-state (IBS) and KING-robust kinship (Manichaikul *et al*. 2010) among the 84 BRBH animals in PLINK using 13,875 LD-pruned SNPs (*r*² < 0.1). We classed pairs as first-, second-, or third-degree relatives at kinship > 0.177, > 0.0884, and > 0.0442. We tested the difference in mean IBS within *high*-BRBH and within *low*-BRBH by permuting group labels 10,000 times.

### Composite selection signals (CSS) analysis

To detect selection signatures across the BRBH genome, we used the multipronged composite selection signal (CSS) developed by Randhawa and colleagues for use in livestock populations (Randhawa *et al*. 2014). The CSS method generates a composite statistic (*CSS score*) for each autosomal SNP from population genetic summary statistics. These statistics were calculated as detailed below using the VCFtools (v.0.1.16) and bcftools (v.1.23.1) software tools (Danecek *et al*. 2011; Danecek *et al*. 2021).

For this analysis, we used the *high*-BRBH samples and all samples from a single comparator production breed: Holstein (HOLS; *n* = 83). We used four population genetic summary statistics to compute *CSS scores* for each SNP. First, within-population nucleotide diversity (*π*) (Nei 1987) was estimated for each population independently in sliding 1 Mb windows with a 500 kb step. Weir and Cockerham’s *F*_ST_ statistic (Weir & Cockerham 1984) was then estimated between the *high*-BRBH and HOLS groups (per-site and in sliding 1 Mb/500 kb windows). Tajima’s *D* statistic (Tajima 1989) was computed per population in non-overlapping 1 Mb windows directly from the per-population VCFs. The directional change in selected allele frequency (*ΔSAF*) (Randhawa *et al*. 2014) was computed in R (v.4.3.2) from per-site allele frequencies generated with VCFtools. For each statistic (*π*, *F*_ST_, *D*, and *ΔSAF*), we converted raw values to fractional ranks as described in Randhawa *et al*. (2014). To reduce single-SNP noise and emphasise the contiguous signal, we smoothed per-SNP *CSS scores* across the genome by averaging within 1 Mb sliding windows advanced in 50 kb steps. We visualised both raw and smoothed *CSS scores* as Manhattan plots, with a significance threshold set at the 99.9^th^ percentile of the empirical *CSS score* distribution. To protect against isolated outlier SNPs driven by genotyping artefacts or single-marker noise, each candidate was required to be supported by at least one additional SNP from the top 1% of the distribution within a ±1 Mb flanking window. We retained only candidates meeting this contiguity criterion as validated selection signals.

To further investigate the functional biology of the selection signatures, gene annotation was performed using the Bovine Genome Database BovineMine (v1.7) integrated data warehouse resource (Kambal *et al*. 2026). We extracted the genomic region spanning ±1.0 Mb around each defined selection signature peak and determined gene content using BovineMine. The current version of BovineMine (v1.7) uses the ARS-UCD2.0 genome assembly build, a minor update to ARS-UCD1.2. To provide information for functional interpretation, the genes identified within the genomic selection signature regions were used for gene set overrepresentation analysis (GSORA) with the g:GOSt tool (https://biit.cs.ut.ee/gprofiler/gost) within the g:Profiler web server (v. 0.2.2; 19^th^ April 2026 release) (Kolberg *et al*. 2023).

## Results

### Results of SNP filtering and quality control, and breed genomic diversity

**Supplementary Table 2** shows the number of SNPs remaining after basic SNP filtering and QC steps. The BRBH breed (*n* = 84) exhibited the highest mean genome-wide SNP *H*_o_ value (0.3697). As noted by Browett *et al*. (2018) the extremely low mean genome-wide SNP *H*_o_ value observed for the West African NDAG population is almost certainly a consequence of ascertainment bias introduced by the focus on polymorphic SNPs in European *B. taurus* during design of the BovineSNP50 BeadChip (Matukumalli *et al*. 2009; McTavish & Hillis 2015).

### Genetic structure of European cattle breeds and identification of a BóRiabhach ancestry component

ADMIXTURE results for *K* = 2 to 9 are shown in **Figure 2A** (*K* = 2–14 in **Supplementary Figure 1**). A component concentrated in the BRBH first appeared at *K* = 9: The BRBH had a mean of 0.37 on this component, no other breed averaged more than 0.07, and no non-BRBH animal exceeded 0.33 (**Supplementary Table 3**). The component and its membership were stable from *K* = 9 to 12 (correlation ≥ 0.998 between *K*); at *K* = 13 it divided, with 11 animals retaining a high value on one of the two resulting components. Using a threshold of 0.50 at *K* = 9, we classified 28 animals as *high*-BRBH and 56 as *low*-BRBH (**Figure 2B**). BóRiabhach ancestry ranged from >0.99 to <0.04, and 25 animals had < 0.10. We used the 28 *high*-BRBH animals as a separate population for some downstream analyses, including *N*_e_-based demographic modelling, analysis of genomic inbreeding using *F*_ROH_ values, and selection signature detection using the CSS method.

**Figure 2:**
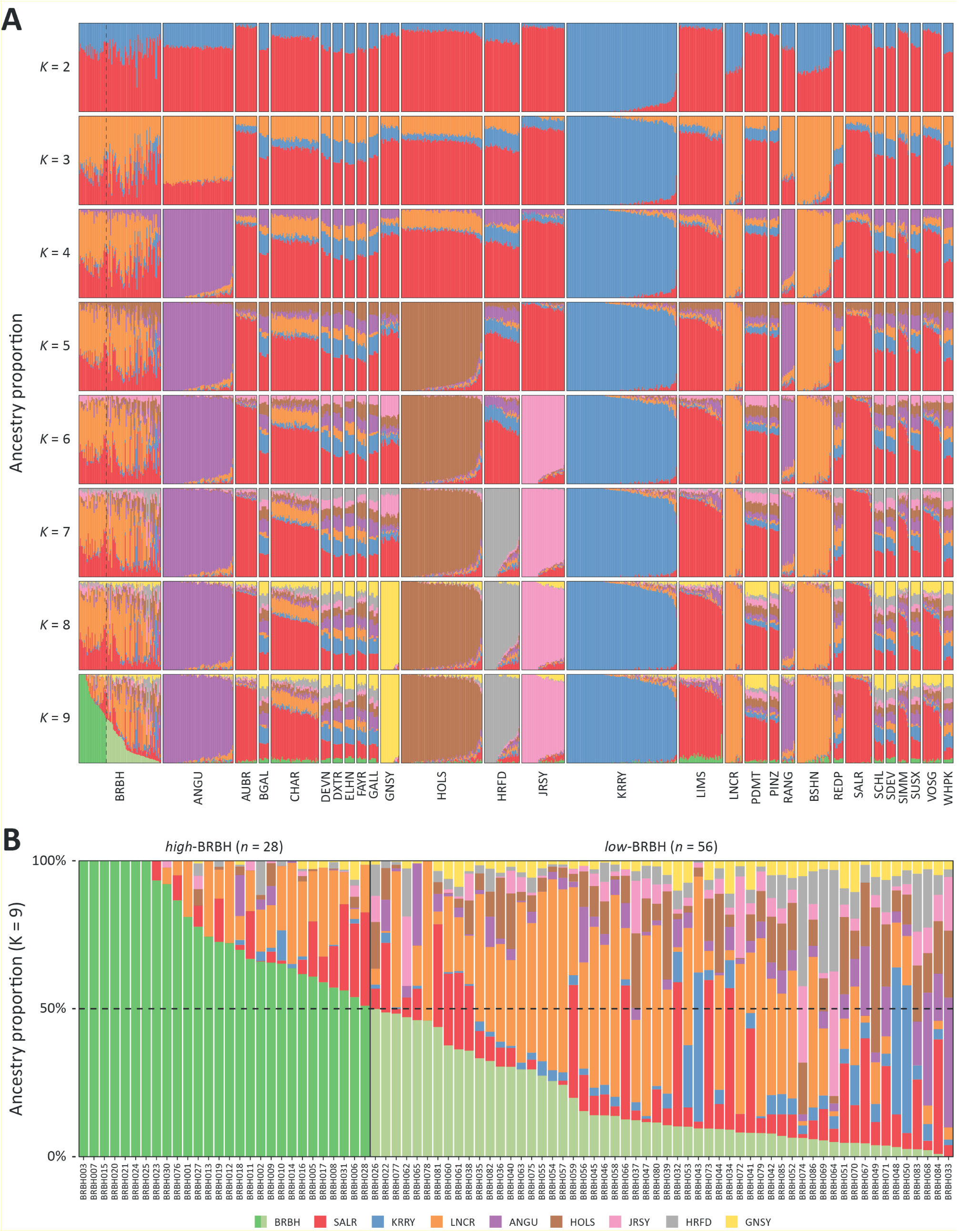
(**A**) ADMIXTURE analysis of individual animals using genome-wide SNP data (13,653 LD-pruned autosomal SNPs) for 772 cattle. Results are shown for separate analyses with modelled ancestral populations *K* = 2 to 9. The results for *K* = 9 modelled ancestral populations revealed a BRBH-diagnostic ancestry component substantially enriched in the BóRiabhach (BRBH) breed sample (*n* = 84). The 28 BRBH animals with a BRBH-diagnostic ancestry component > 0.50 (50%) (*high*-BRBH) are outlined with a vertical line and a dashed horizontal line, and the remaining 56 animals with a BRBH ancestry component < 0.50 (50%) are indicated in lighter green (*low*-BRBH). (**B**) Results from the ADMIXTURE analysis (*K* = 9) (13,653 autosomal SNPs). The 28 BRBH animals with a BRBH-diagnostic ancestry component > 0.50 (50%) are delineated with a line showing the cut-off (*high*-BRBH); the remaining 56 animals with a BRBH ancestry component < 0.50 (50%) follow those (*low*-BRBH). The individual BRBH animal sample codes are indicated below each ancestry bar.

### Multivariate principal component analysis (PCA) of genome-wide SNP data in 29 European cattle breeds

**Figure 3** shows the PCA results for 13,653 autosomal SNPs from 772 animals representing 29 European cattle breeds. As with the structure plot, PC1 broadly separates the Kerry (KRRY), British Shorthorn (BSHN), and Lincoln Red (LNCR) from other European breeds, with Jersey (JRSY) samples clustered at the opposite end of the PC1 axis. The second PC (PC2) separates KRRY from BSHN and LNCR, with the other European breeds arrayed at intermediate positions along the PC2 axis.

**Figure 3:**
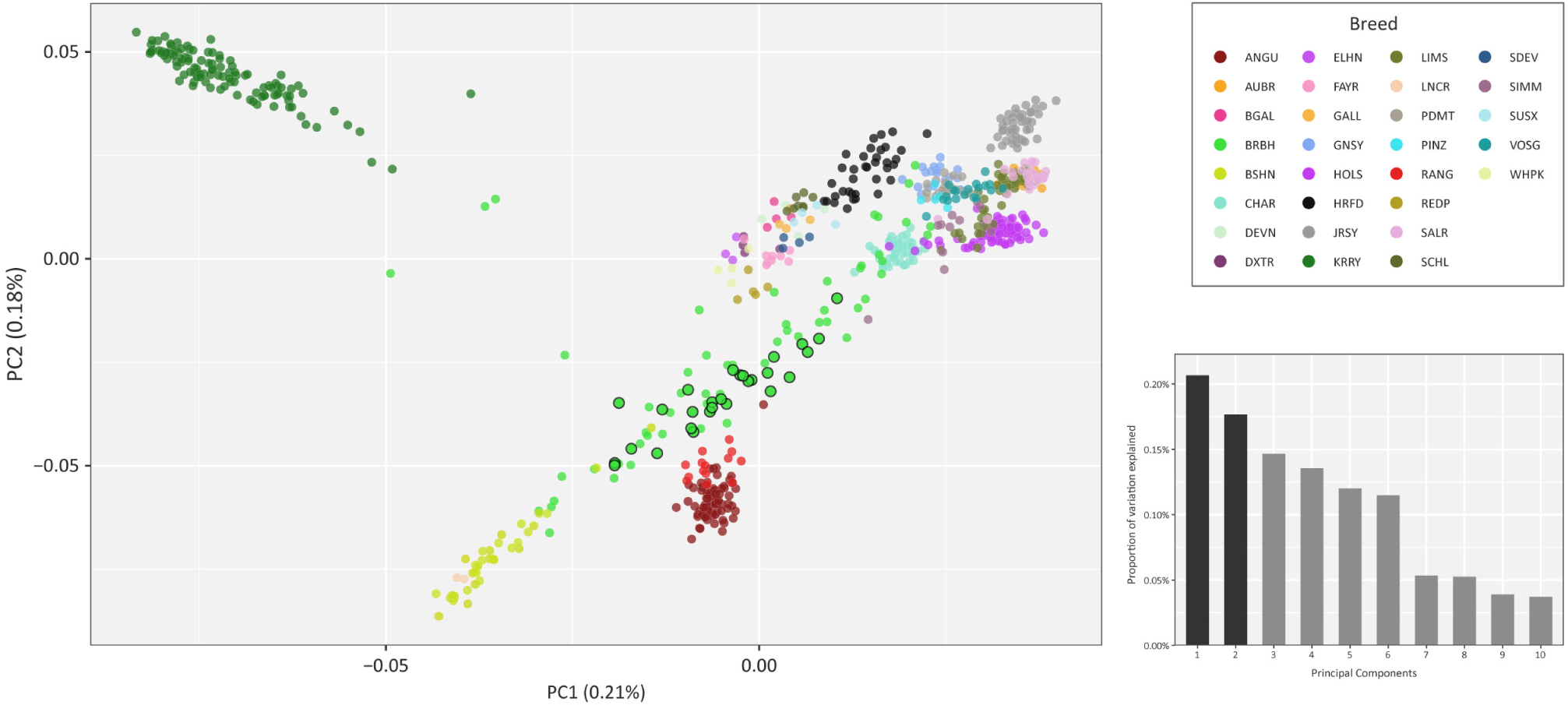
Principal component analysis (PCA) plot of PC1 and PC2 from genome-wide SNP data (13,653 autosomal SNPs) for 772 animals representing 29 European cattle breeds. The smaller histogram shows the variance contributions of the first 10 PCs relative to the total variance across all 772 PCs. PC1 and PC2 account for 0.21% and 0.18% of the total variation, respectively. The data points for the 28 BóRiabhach (BRBH) animals with a BRBH-diagnostic ancestry component > 0.50 (50%) are indicated with a dark border (*high*-BRBH).

The 84 BRBH animals are shown as bright green circles on the PCA, and the *high*-BRBH group animals identified from the BRBH ancestry component in **Figure 2B** are indicated with dark borders. The most notable feature of the total BRBH breed sample is that it forms a broad swathe across both PC1 and PC2, with the *high*-BRBH group more clustered across PC2 and, to a lesser extent, across PC1. The marked lack of clustering in the 56 *low*-BRBH, compared to the other European breeds, suggests substantial genetic heterogeneity due to crossbreeding and gene flow from other cattle populations.

### Phylogenetic ancestry graphs of taurine cattle breeds

We used TreeMix to model population splits and gene flow between *high*-BRBH and 29 comparator breeds (**Figure 4**). The proportion of variance in relatedness explained rose from 0.909 with no migration to 0.939 with two edges; each additional edge added < 1%, so we present the *m* = 2 graph. After the NDAG root, the Piedmontese (PDMT) and then a clade of seven continental breeds (Aubrac, AUBR; Charolais, CHAR; Limousin, LIMS; Pinzgauer, PINZ; Salers, SALR; Simmental, SIMM; and Vosgienne, VOSG) split from the remaining breeds. The *high*-BRBH was a sister group to BSHN and LNCR, and this clade was sister to Angus (ANGU) and Red Angus (RANG). The stronger migration edge ran from LIMS into *high*-BRBH (weight = 0.28); it was recovered in all 10 replicates at every *m* value from 2 to 8 (weights 0.28–0.32). The second edge ran from the branch ancestral to *high*-BRBH, BSHN and LNCR into CHAR (weight = 0.27) and was likewise recovered at every *m*. From *m* = 4, an additional edge from this branch into HOLS was inferred (weight ≈ 0.21). No edge from HOLS into *high*-BRBH was inferred at any *m*. None of the *f*_3_ statistics with *high*-BRBH as target were negative (all *Z* > 15), as expected when strong drift in the target population masks the admixture signal.

**Figure 4:**
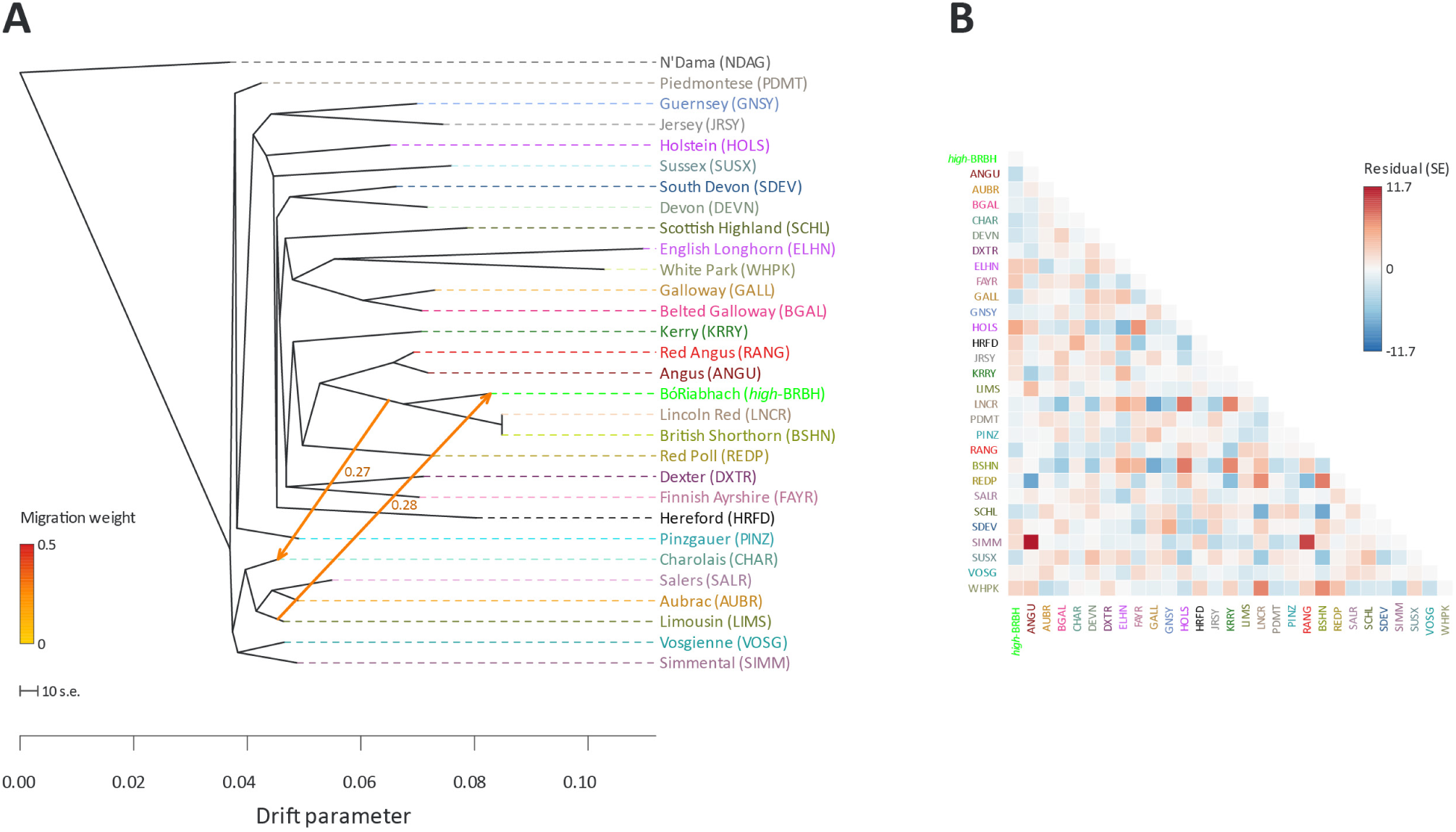
Phylogenetic tree network graph with two migration edges (*m* = 2) derived from genome-wide SNP data for 29 European cattle breeds. The analysis used 28 *high*-BRBH animals and 28 European comparator breeds, plus the West African taurine N’Dama (NDAG) population (738 animals; 25,395 autosomal SNPs). Migration-edge weights are shown beside each arrow. The NDAG is included as an outgroup population. Coloured lines and arrows indicate migration edges that model gene flow between lineages, with migration weights represented by the colour gradient.

### Modelling recent and historical effective population size (*N*_e_) using genome-wide SNP data

We estimated historical effective population size (*N*_e_) trajectories for *high*-BRBH and all European comparator breeds (**Supplementary Table 4**). **Figure 5** shows *high*-BRBH alongside three production breeds (HOLS, JRSY and RANG) and four traditional heritage breeds (Belted Galloway, BGAL; Dexter, DXTR; KRRY; and LNCR). The most recent estimate corresponds to 13 generations before sampling (approximately 78 years), or the 1920s–1940s for breeds sampled between 2000 and 2020. Because estimates depended strongly on sample size when *n* < 20 (**Supplementary Figures 3 and 4**), we restricted comparisons to breeds with *n* > 20. Among these, *high*-BRBH had the lowest recent *N*_e_ (64), and only KRRY (*n* = 113; *N*_e_ = 92) was also below 100; all other breeds with *n* > 20 had *N*_e_ ≥ 111. *N*_e_ for *high*-BRBH declined steadily from approximately 1,800 at 900 generations ago to 64, a trajectory most similar to that of KRRY (**Figure 5**). However, we could estimate *N*_e_ for KRRY only back to 552 generations because fewer SNP pairs were informative in that breed.

**Figure 5:**
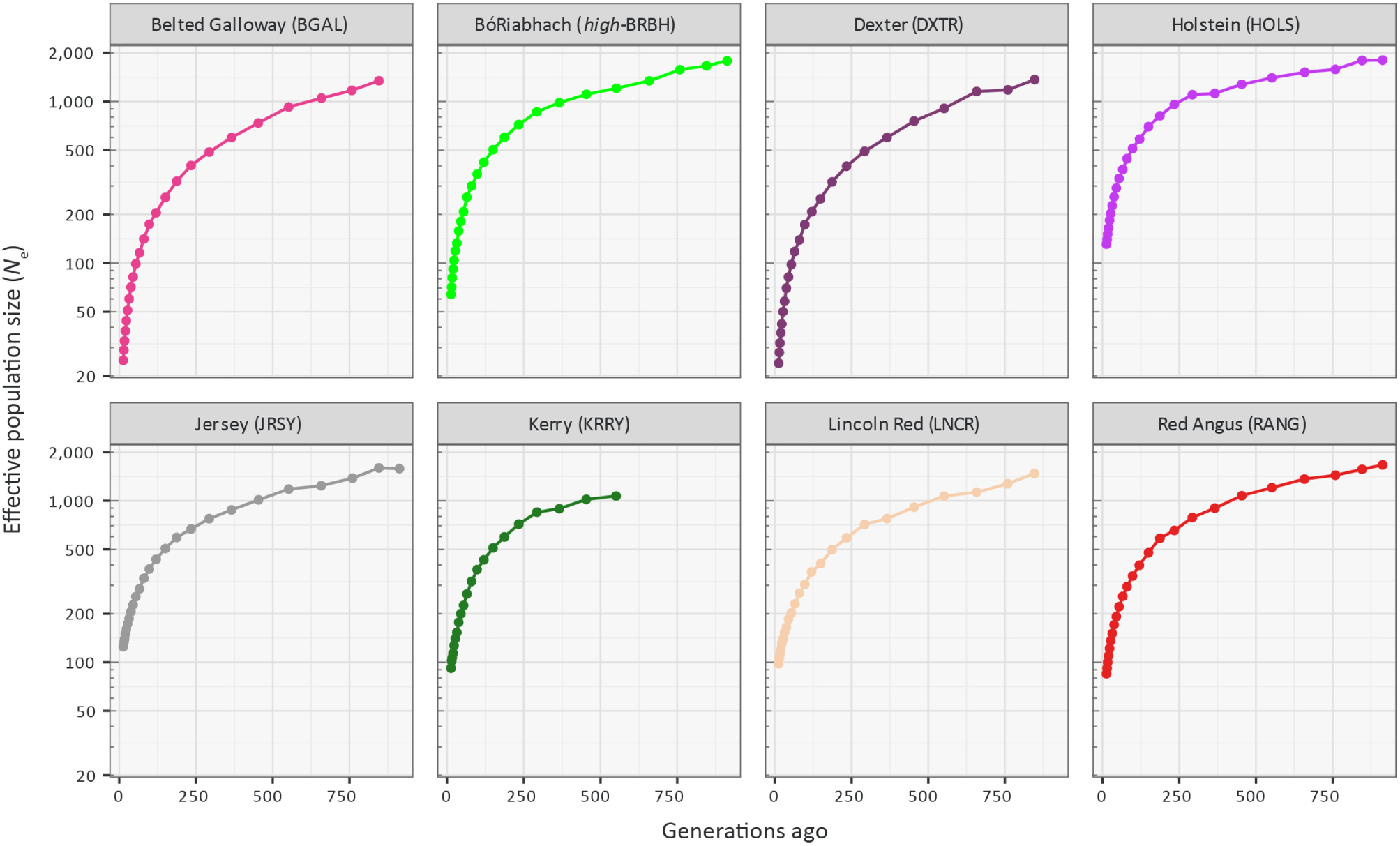
Modelling effective population size (*N*_e_) for the *high*-BRBH BóRiabhach breed sample and seven comparator breeds, comprising production and traditional heritage cattle breeds. Effective population size (*N*_e_) is shown on a logarithmic scale.

### Detection of runs of homozygosity (ROH) and estimation of *F*_ROH_

We identified runs of homozygosity (ROH) in 764 animals using 25,905 autosomal SNPs (**Supplementary Tables 5-6**; **Figure 6**). Mean *F*_ROH_ ranged from 0.006 (PDMT) to 0.224 (White Park, WHPK). The highest values were in British and Irish heritage breeds (WHPK; English Longhorn, ELHN; and DXTR; all ≥ 0.19), where most ROH length was in segments > 8 Mb (**Figure 6A**). Among production breeds, *F*_ROH_ was highest in Hereford (HRFD, 0.109) and lowest in the continental beef breeds CHAR, AUBR, LIMS and SIMM (0.020–0.023).

**Figure 6:**
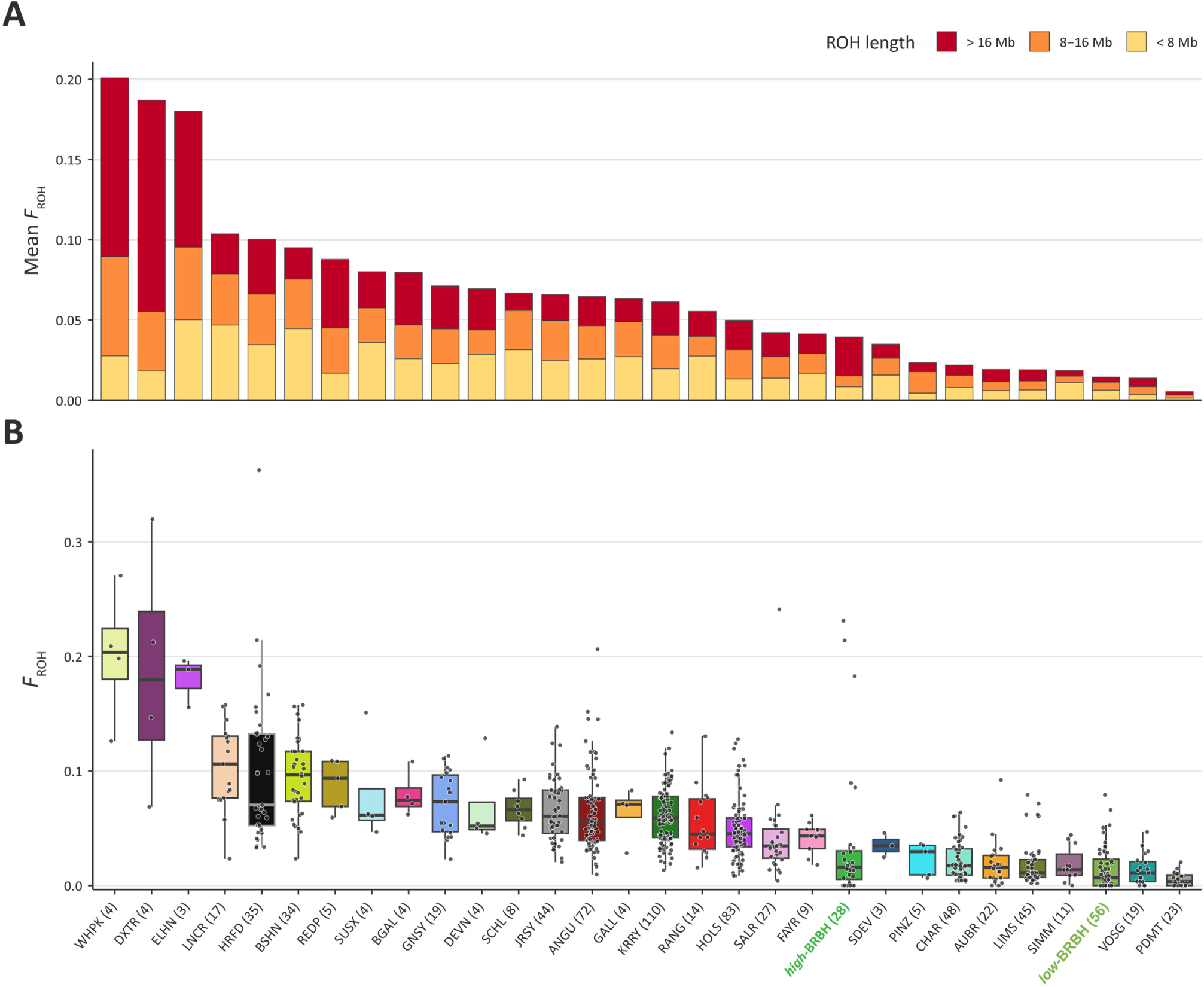
(**A**) Stacked bars showing mean *F*_ROH_ values for the *high*-BRBH and *low*-BRBH BóRiabhach population samples alongside 28 comparator breeds. The figure also shows the proportions of summed ROH, partitioned into defined length categories. (**B**) Box plots showing individual *F*_ROH_ values for all animals across the *high-* BRBH and *low-*BRBH population samples and the 28 comparator breeds. The median for each breed/population sample is indicated by a thick horizontal line; the boxes show the interquartile ranges (IQRs), and the whiskers extend to 1.5 × IQR for each breed/population sample, with outliers also evident.

The *low*-BRBH had the third-lowest mean *F*_ROH_ (0.016), above only Vosgienne (VOSG, 0.014) and PDMT (0.006), and no ROH were detected in 20 of the 56 *low*-BRBH animals. Mean *F*_ROH_ in *high*-BRBH (0.043) was comparable to SALR and Finnish Ayrshire (FAYR) (both 0.044) but concealed a highly uneven distribution (**Figure 6B**): the median was 0.017, six animals had no detected ROH, and three animals had values exceeding the highest breed means (BRBH020, 0.265; BRBH025, 0.231; and BRBH015, 0.196), almost entirely in ROH > 8 Mb and mostly > 16 Mb.

### Genomic relatedness among animals sampled from the BóRiabhach breed

We estimated pairwise IBS and KING-robust kinship for the 84 BRBH animals using 13,875 LD-pruned SNPs (**Figure 7**; **Supplementary Table 7**). Mean IBS was higher within the *high-*BRBH (0.737) than within the *low-*BRBH (0.696) or between the groups (0.698), and this difference was greater than in any of 10,000 random relabellings of the animals (*P* < 0.0001). Inspection of the kinship results showed why: within the *high-*BRBH, 10.1% of pairs were first-degree relatives and a further 24.1% second-degree, compared with 1.2% and 2.5% within the *low-*BRBH; in addition, fewer than half of *high-*BRBH pairs were unrelated. Every *high-*BRBH animal had at least one first-degree relative in the core, and three animals (BRBH003, BRBH076, and BRBH023) each had 9–13 first-degree relatives among the other 27, consistent with extensive use of a small number of sires. The most inbred animals, BRBH015 and BRBH020, were themselves first-degree relatives, and both were first-degree relatives of BRBH023.

**Figure 7:**
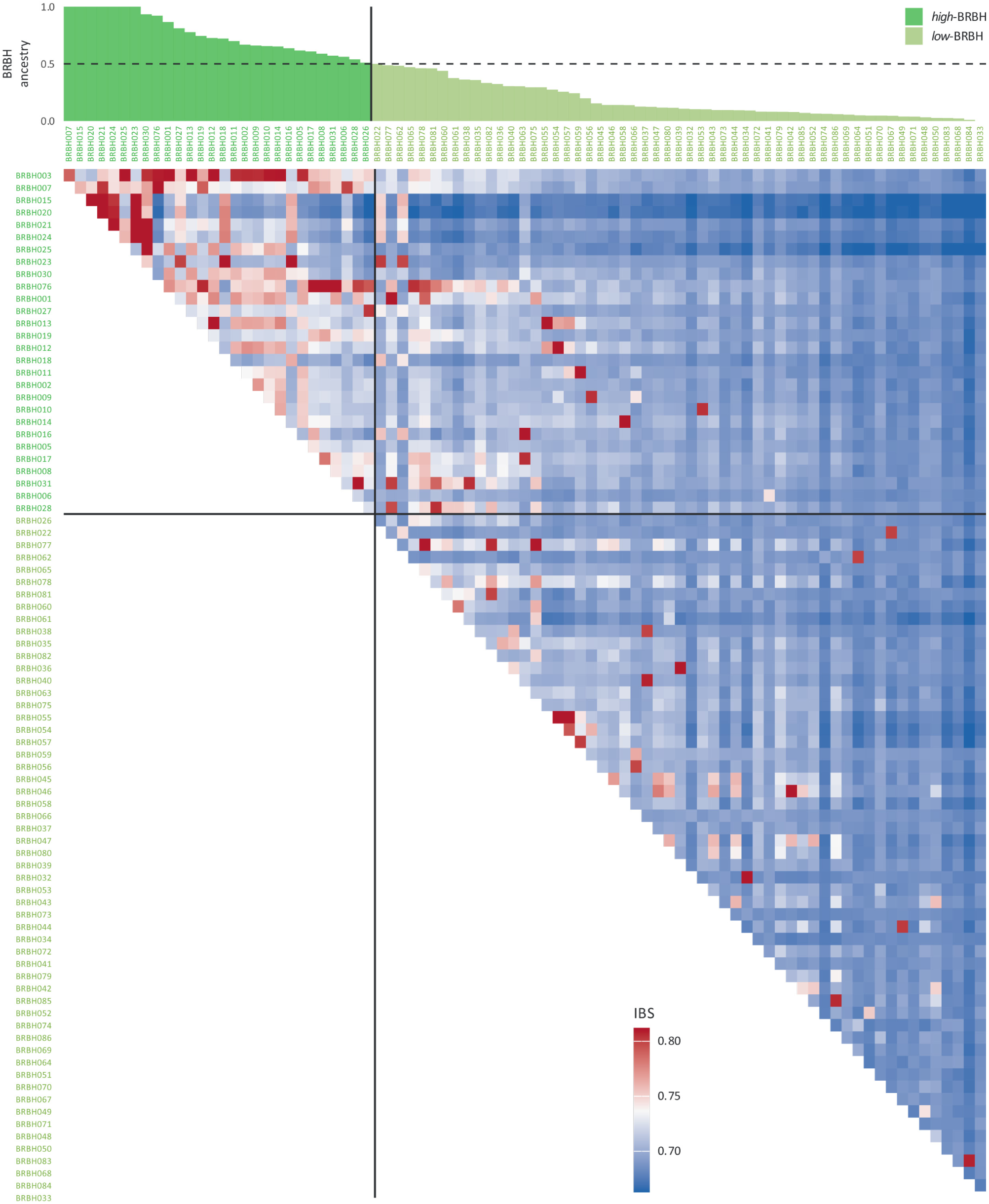
Upper-triangular pairwise identity-by-state (IBS) matrix for all BóRiabhach (BRBH) animals (*n* = 84), ordered by BRBH ancestry; animal codes are shown in green for *high*-BRBH (*n* = 28) and light green for *low*-BRBH (*n* = 56), with horizontal and vertical lines marking the boundary between the two groups. The bar chart above the matrix shows each animal’s BRBH ancestry component at *K* = 9, with the 0.50 threshold indicated.

### Detection of selection signatures across the BóRiabhach genome

We used 24,380 genome-wide SNPs for the CSS analyses in the *high*-BRBH (*n* = 28) and HOLS (*n* = 83) groups. We used these 24,380 SNPs because they produced results for all four selection statistics used in the CSS analyses (*π*, *F*_ST_, Tajima’s *D*, and *ΔSAF*). The results of the CSS analysis are shown in unsmoothed form in **Figure 8A** and in smoothed form in **Figure 8B**, and **Supplementary Table 8** shows the focal SNPs within the *CSS score* peaks defined as selection signals for the unsmoothed analysis (top 0.1% and at least one SNP in the top 1.0% within ±1 Mb), and **Supplementary Table 9** shows the focal selection genomic windows identified from the smoothed analysis (1 Mb sliding windows in the top 0.1% of smoothed CSS scores, with overlapping windows merged). **Supplementary Tables 10** and **11** show the genes within the genomic selection signature regions identified from the unsmoothed and smoothed CSS analyses, respectively.

**Figure 8:**
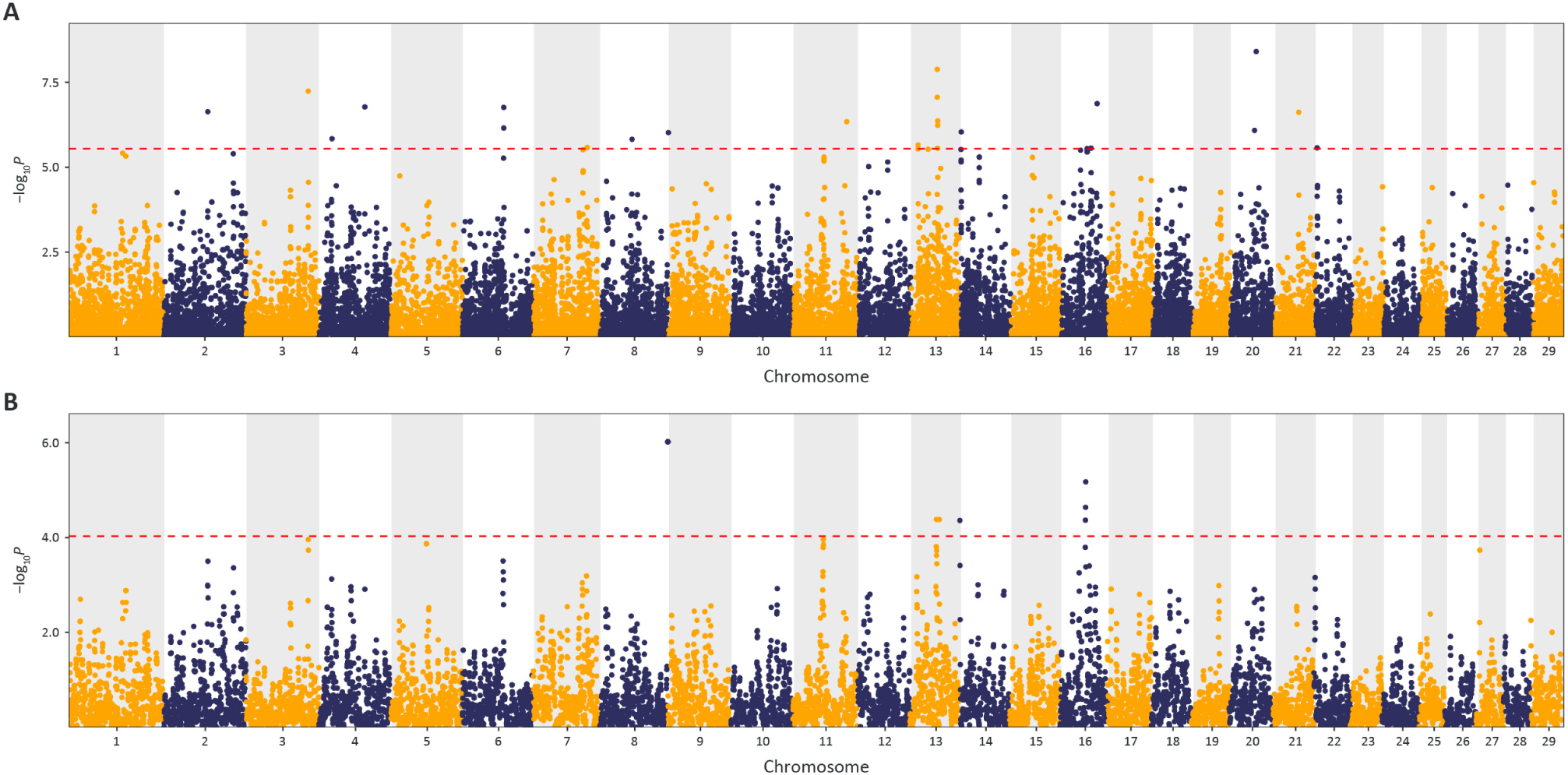
Manhattan plots of composite selection signal (CSS) results for the *high*-BRBH BóRiabhach population sample (*n* = 28) compared with the Holstein (HOLS) breed sample (*n* = 83). (**A**) Unsmoothed results. (**B**) Smoothed results obtained by averaging *CSS scores* across SNPs within each 1 Mb window. The red dotted line on each plot denotes the genome-wide 0.1% threshold for the empirical *CSS scores*.

The gene sets identified from the unsmoothed (324 genes) and smoothed (177 genes) CSS analyses were used for functional gene set overrepresentation analysis (GSORA) with the g:GOSt tool within g:Profiler, and the results for the smoothed CSS analysis (see **Figure 8B**) are shown in **Figure 9** and detailed in **Supplementary Table 12**. In addition, **Supplementary Figure 2** and **Supplementary Table 13** show the equivalent results for the unsmoothed CSS analysis. The g:SCS *P*-value adjustment method specifically developed for multiple test correction with g:GOSt was used, which is more conservative than the Benjamini-Hochberg FDR method but less strict than the Bonferroni correction (Raudvere *et al*. 2019).

**Figure 9:**
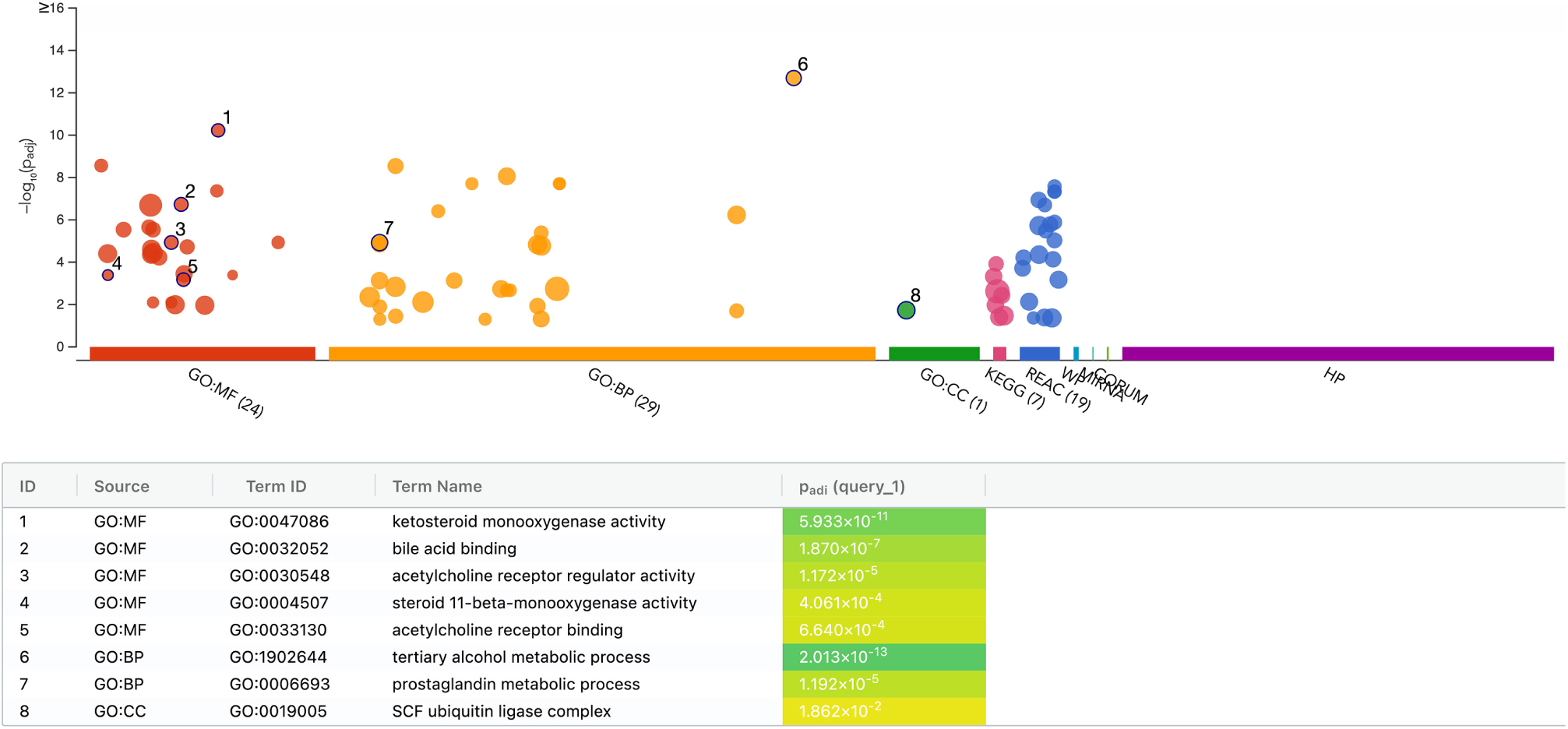
Manhattan plot of the g:GOSt GSORA outputs from the smoothed CSS analysis results (177 genes). Only significant functional terms are shown (g:SCS *P*_adj._ < 0.05), and the functional entities are from the Gene Ontology (GO), KEGG, Reactome, WikiPathways (WP), miRTarBase, CORUM, and Human Phenotype Ontology (HP). The GO driver terms are numbered and shown with the g:SCS *P*_adj._ values for these GO terms (for further methodological details see: Kolberg *et al*. 2023).

Functional overrepresentation analysis of the 177 genes within the significant regions from the smoothed CSS analysis identified 80 significant terms: 24 enriched Gene Ontology (GO) molecular function terms, 29 biological process terms, and one cellular component term, along with seven KEGG and 19 Reactome pathways (**Figure 9; Supplementary Table 12**). Eight GO driver terms were highlighted. The five significant molecular function driver terms were *GO:0047086, ketosteroid monooxygenase activity* (*P*_adj._ = 5.933 × 10^−11^); *GO:0032052, bile acid binding* (*P*_adj._ = 1.870 × 10^−7^); *GO:0030548, acetylcholine receptor regulator activity* (*P*_adj._ = 1.172 × 10^−5^); *GO:0004507, steroid 11-beta-monooxygenase activity* (*P*_adj._ = 4.061 × 10^−4^); and *GO:0033130, acetylcholine receptor binding* (*P*_adj._ = 6.640 × 10^−4^). The two significant biological process driver terms were *GO:1902644, tertiary alcohol metabolic process* (*P*_adj._ = 2.013 × 10^−13^), the most significantly enriched term overall, and *GO:0006693, prostaglandin metabolic process* (*P*_adj._ = 1.192 × 10^−5^). The single significant cellular component driver term was *GO:0019005, SCF ubiquitin ligase complex* (*P*_adj._ = 1.862 × 10^−2^). The unsmoothed CSS gene set (324 genes) gave a closely similar profile, with 68 significant terms and seven of the same eight GO driver terms, lacking only the cellular component term **(Supplementary Figure 2; Supplementary Table 13**).

## Discussion

This study is the first genomic characterisation of the Irish traditional BóRiabhach cattle breed, and four findings stand out. First, the breed belongs to a British-Irish taurine lineage but is distinct from the other Irish breeds sampled. Second, a distinctive BRBH ancestry component is concentrated in a minority of animals, while the remainder show extensive admixture, chiefly from Limousin and Holstein. Third, the high-ancestry animals in the core group are closely related, with recent inbreeding evident in a few individuals. Fourth, much of the breed’s apparent genetic diversity reflects recent crossbreeding rather than retained native variation. Taken together, these results present a central challenge for conserving the BóRiabhach breed: the animals that best represent the breed are also the least genetically diverse.

### Origins and relationships

In the TreeMix phylogenetic analysis, the *high*-BRBH (BRBH-diagnostic ancestry > 0.50) was a sister group to the British Shorthorn and Lincoln Red, within a clade that also included Angus and Red Angus (**Figure 4**). This is unsurprising and consistent with 19^th^-century accounts linking the brindled Irish cattle to the unimproved native British cattle and Ireland (Wilde 1857). This is also the first molecular evidence for understanding the breed’s deeper origins. The *high*-BRBH did not group with the other two traditional Irish breeds in the panel, the Kerry and Dexter. This may, in part, be due to genetic drift in the Kerry, a population that has long been closed off and has a small effective population size (Browett *et al*. 2018). Even so, this suggests that the BóRiabhach is not merely an offshoot of the Kerry lineage. Importantly, however, the finer-scale relationships within this clade and across the phylogenetic analysis should be interpreted cautiously, as several British and Irish comparator breeds were represented by five or fewer animals (BGAL; DEVN; DXTR; ELHN; GALL; Red Polled, REDP; South Devon, SDEV; Sussex, SUSX; and WHPK).

TreeMix identified Limousin as the clearest source of gene flow into the high-ancestry BRBH core group, with this edge recovered in every replicate at every model size (**Figure 4**). *Low*-BRBH animals were distributed across PC1 and PC2 rather than forming a single intermediate cluster (**Figure 3**). This pattern points to repeated crossing with several breeds over multiple generations rather than a single admixture event (McVean 2009).

### Defining the BóRiabhach genomically

Independent of pedigree, which has not been systematically recorded for this breed, the BRBH-diagnostic ancestry component identified at *K* = 9 in the ADMIXTURE analysis (see **Figure 2**) provides a basis for defining breed membership. Animals registered as BóRiabhach span almost the full range of this component, from >98% to <4%, and more than a quarter carry less than 10% (**Figure 2B**; **Supplementary Table 3**). However, the 0.50 (50%) cut-off we used to define *high*-BRBH animals is arbitrary, and any working classification the BoRiabach Cattle Society adopts should account for how sensitive downstream results are to alternative cut-off values.

In addition, caveats apply to this BRBH-diagnostic ancestry component. First, it is defined relative to this comparator panel and without historical reference samples. Therefore, it captures what distinguishes the present-day core animals from other breeds, not necessarily the population’s ancestry before its 20^th^-century decline. Second, ADMIXTURE components can form around clusters of closely related individuals, particularly in the most heavily sampled population (Lawson *et al*. 2018). Relatedness within the core is high (**Figure 7**), and the most inbred animals are among those with the highest BRBH ancestry. Taken together, these observations suggest that some of this component may reflect a small number of closely related families.

Substructure within the population also explains why the pooled BóRiabhach sample had the highest observed heterozygosity among the 30 breeds (**Supplementary Table 1**). This does not indicate good genetic health but rather reflects gene flow from divergent sources, a pattern observed in the revived Belgian Campine and in Slovenian Cika cattle (Simčič *et al*. 2015; Francois *et al*. 2017). Both of these comparisons show why the definition matters. In the Cika, the native genetic background was recovered from admixed animals using haplotype, phenotype and pedigree information. In the Canadienne, recent *N*_e_ estimates ranged from 13 to 35, depending on the purity classification applied (Carrier *et al*. 2023). Similarly, the same dependence on how the breed is defined would apply to the BóRiabhach.

### Genetic health of the high ancestry core

The *N*_e_, ROH and IBS results indicate a small, closely related core population, with evidence for recent inbreeding. Among breeds with sufficiently large sample sizes (*n* > 20), the *high*-BRBH had the lowest recent *N*_e_. Mean relatedness within the *high*-BRBH core was also higher than among *low*-BRBH animals (**Figure 7**; **Supplementary Table 7**).

Mean *F*_ROH_ in the *high*-BRBH group was moderate and lower than in several large production breeds (**Figure 6**). However, this average masks a skewed distribution. Three animals carry most of the long ROH, while six carry no detectable ROH. *F*_ROH_ values of 0.20–0.26, composed almost entirely of ROH > 16 Mb, are consistent with matings between close relatives within the last two to three generations. Two of these animals (BRBH015 and BRBH020) are first-degree relatives of each other, and both are first-degree relatives of BRBH023, one of the most connected animals in the sample. Recent inbreeding has therefore been concentrated in a few lines where BRBH ancestry has accumulated through matings between relatives.

The *N*_e_ estimate of 64 animals for the *high*-BRBH group should be evaluated with particular care (**Figure 5** and **Supplementary Table 4**). The most recent estimate, derived using the SNeP software, refers to approximately 13 generations before sampling (broadly the mid-20^th^ century) and therefore predates the breed’s decline to an estimated 20 animals by 2016. Contemporary *N*_e_ is almost certainly well below 64, and the fact that this historical estimate exceeds the FAO minimum of 50 (Food and Agriculture Organization 2013) should not be taken as evidence that the present population does so. Two further factors affect the estimate. LD-based methods are sensitive to admixture (Waples 2024) and are biased downward when a sample contains close relatives, because relatives share long haplotypes and inflate LD at every distance. The *high*-BRBH sample includes many first-degree relatives and three animals with nine or more first-degree relatives each, so the estimate reflects the sampled core rather than the breed as a whole, and part of its low value likely reflects this family structure rather than the historical population alone. Even taken at face value, it falls well short of the *N*_e_ ≥ 100 recommended for long-term genetic sustainability (Meuwissen 2009), and the related low *N*_e_ of the Kerry, the only other Irish breed with an adequate sample, shows that this is a shared feature of Ireland’s native cattle.

### Selection signatures

The CSS scan identified several candidate regions in *high*-BRBH relative to Holstein (**Figure 8**; **Supplementary Tables 8**–**11**). These results should be treated as exploratory. The scan used 28 animals and 24,380 SNPs. Admixture within the core, including gene flow from Holstein itself, can also distort each component statistic and generate spurious signals (Vatsiou *et al*. 2016).

Functional enrichment of the 177 genes within the smoothed CSS regions returned 80 significant terms (24 GO molecular function, 29 biological process, one cellular component, seven KEGG and 19 Reactome; **Figure 9; Supplementary Table 12**), dominated by steroid, bile acid and prostanoid metabolism, with a smaller group of acetylcholine-receptor terms. This apparent breadth, however, is almost entirely attributable to two loci. A single set of seven paralogous aldo-keto reductase genes, consistent with the aldo-keto reductase family 1 C (AKR1C) gene cluster on BTA13, accounts for the four most significant GO driver terms (ketosteroid monooxygenase activity, bile acid binding, tertiary alcohol and prostaglandin metabolic process) and all 19 Reactome pathways. A second group of genes from the proximal BTA14 region accounts for the remaining steroidogenic terms (three *CYP11B*-like genes constitute the entire GO category for steroid 11β-monooxygenase activity) and, through a neighbouring LY6/uPAR-family cluster, for all of the acetylcholine-receptor terms. Both are tandemly arrayed gene families, in which one candidate region contributes many paralogs to the same small GO categories and inflates over-representation statistics; the enrichment therefore reflects the gene content of two regions rather than a genome-wide functional pattern, and should not be interpreted as evidence of selection on steroid metabolism or cholinergic signalling without independent support.

A selection scan with adequate power will require high-density or whole-genome sequence data from a larger cohort of animals with very high BRBH ancestry. Using this approach with the CSS method, we have previously identified selection signals associated with known breed traits, including heat tolerance, in Latin American Criollo cattle (Ward *et al*. 2024).

### Implications for conservation management

The central management challenge is that preserving distinctiveness and maintaining diversity pull in opposite directions. Breeding only from the highest-ancestry animals would prioritise reproduction in a small, related group and accelerate inbreeding, while the unrestricted use of admixed animals would continue to erode the BRBH component.

From these results, we can draw several practical steps. First, all extant animals should be genotyped and BRBH ancestry recorded at registration, potentially within graded herdbook sections. Second, mating decisions should weigh genomic relatedness alongside ancestry, for example through optimal contribution selection, so that BRBH ancestry can be increased while the rate of inbreeding is limited (Meuwissen 2009). Put into practice, this means avoiding heavy use of the few very-high-ancestry animals that are already quite closely related. It also means using intermediate-ancestry animals in structured grading-up rather than excluding them. Finally, semen and embryos from core animals should be cryopreserved as a safeguard, in line with the FAO Global Plan of Action for Animal Genetic Resources (Food and Agriculture Organization 2007).

### Limitations and future work

Most British heritage comparators were represented by five or fewer animals. This limits the resolution of the phylogenetic analysis and renders their *N*_e_ estimates unreliable (see **Supplementary Figures 3 and 4**). The medium-density array constrains both the detection of short ROH and the power of the selection scan. The absence of systematic pedigree records means that genomic and genealogical estimates of inbreeding cannot yet be compared.

Most importantly, without historical reference samples, it is not possible to determine how closely the BRBH component reflects the breed as it existed before its twentieth-century decline. Whole-genome sequencing of a larger high-ancestry cohort (*n* > 20 with > 90% BRBH ancestry) would strengthen the selection scan analysis. Combined with genotyping of historical material where available, it would also help address the question of breed authenticity.

### Conclusions

This study presents the first genomic characterisation of the BóRiabhach traditional heritage cattle breed. The breed belongs to a British-Irish taurine lineage and carries a distinctive ancestry component. However, this ancestry is concentrated in a minority of animals (28 of 84), while the remainder show extensive introgression, primarily from Limousin and Holstein. Much of the breed’s apparent genetic diversity therefore reflects crossbreeding rather than retention of native variation. This high-ancestry core is small, closely related, and, in some lines, recently inbred. The estimated historical effective population size of 64 predates the breed’s decline to around 20 animals, so the contemporary effective population size is likely to be substantially lower. Conserving the BóRiabhach will therefore require balancing breed distinctiveness with genetic diversity.

We therefore recommend the following steps for the BóRiabhach Cattle Society: 1) genotype all extant animals; 2) record genomic ancestry at registration; 3) manage matings based on genomic relatedness as well as ancestry; and 4) cryopreserve germplasm from core animals. The results presented here provide a genomic baseline for such a programme.

## Author Contributions

**Darren McLoughlin:** conceptualisation, formal analysis, funding acquisition, investigation, visualisation, writing – original draft, and writing – review & editing. **Gillian P. McHugo:** conceptualisation, formal analysis, investigation, methodology, visualisation, writing – review & editing. **Paul Flynn:** data curation, resources, and writing – review & editing. **Patrick Sullivan:** conceptualisation, funding acquisition, resources, and writing – review & editing. **Sam S. Browett:** conceptualisation, data curation, resources, and writing – review & editing. **David E. MacHugh:** conceptualisation, investigation, project administration, resources, supervision, writing – original draft, and writing – review & editing. **James A. Ward:** conceptualisation, formal analysis, investigation, methodology, resources, software, supervision, validation, visualisation, writing – original draft, and writing – review & editing.

## Acknowledgments

We thank Jens Carlsson, Nettan Carlsson and Hans Visser for useful advice and guidance. We also thank Trustees of the Irish Rare Breeds Society, Matthew English Hayden, Mark McConnell, Patrick McConnell, Sean Carolan, Sean Hyland, and Michael O’Halloran. We also acknowledge support and help with data provision from members of the BóRiabhach Cattle Society.

## Ethics Statement

Weatherbys Scientific (Co. Kildare, Ireland) purified DNA samples for genotyping from hair follicles and nasal swabs. The owners of the cattle provided these biological samples for routine genetic improvement and population monitoring. The European Union (Protection of Animals Used for Scientific Purposes) Regulations 2012 (S.I. No. 543 of 2012) do not classify the collection of hair follicles or nasal swabs as an invasive procedure.

## Data Availability Statement

The Illumina^®^ BovineSNP50 BeadChip dataset for the 84 BóRiabhach cattle used in this study is available from the Dryad Digital Repository (http://doi.org/10.5061/dryad.zs7h44jt8). The equivalent SNP dataset for the 111 Kerry cattle published by Browett *et al*. (2018) is also available from Dryad (https://doi.org/10.5061/dryad.8fk81). The SNP data from the remaining 28 breeds (27 European and one African) are available from the *WIDDE* resource (Sempéré *et al*. 2015).

