## Supplementary Materials for "Genomic Characterisation of the Irish Traditional BóRiabhach Cattle Breed"

### Supplementary Figures

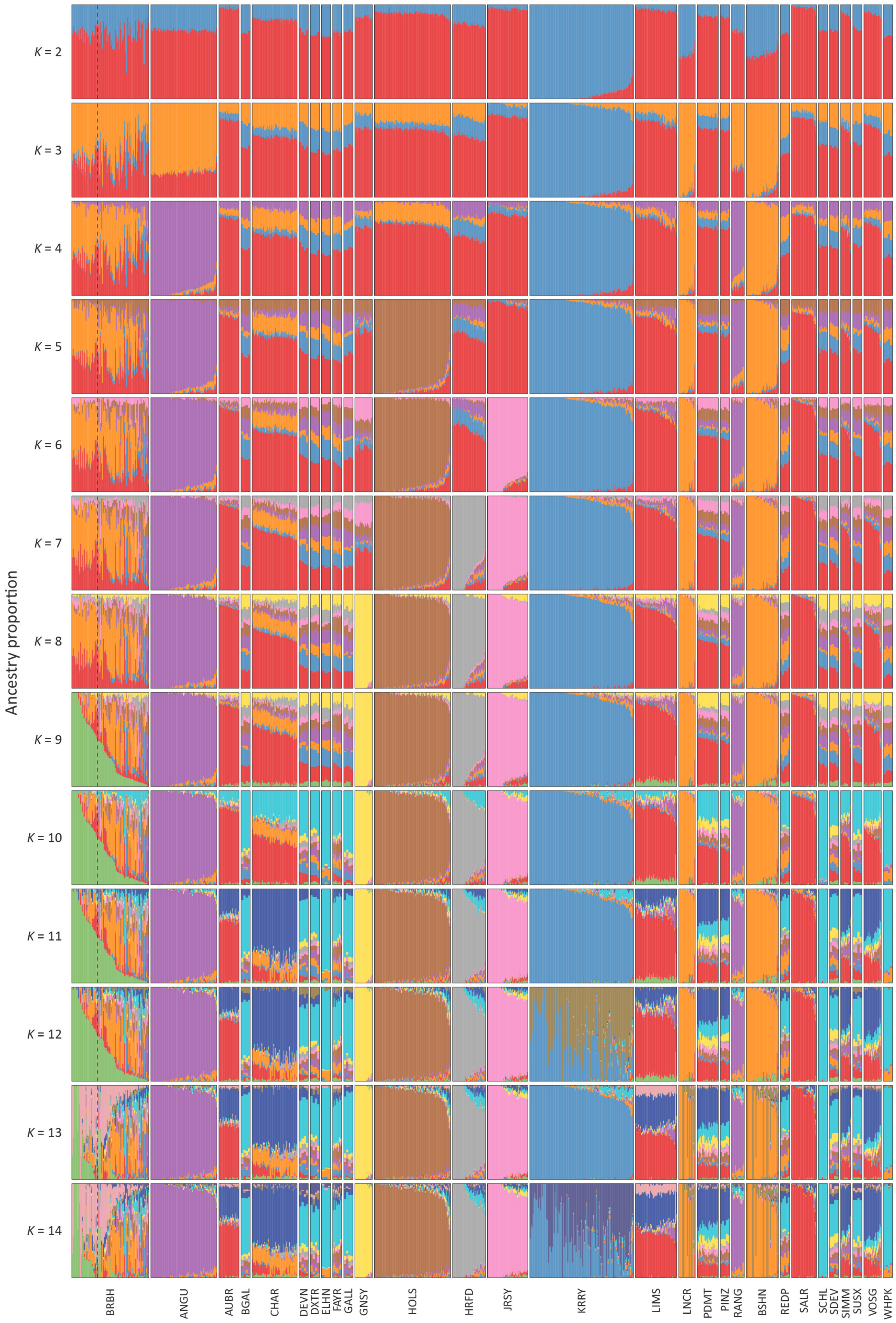


**Supplementary Figure 1:** ADMIXTURE analysis of individual animals using genome-wide SNP data (13,653 LD-pruned autosomal SNPs) for 772 cattle. Results are shown for separate analyses with modelled ancestral populations *K* = 2 to 14. The results for *K* = 9 modelled ancestral populations revealed a BRBH-diagnostic ancestry component enriched in the BóRiabhach (BRBH) breed sample (*n* = 84). The 28 BRBH animals with a BRBH-diagnostic ancestry component > 0.50 (50%) (*high*-BRBH) are outlined with a vertical line.

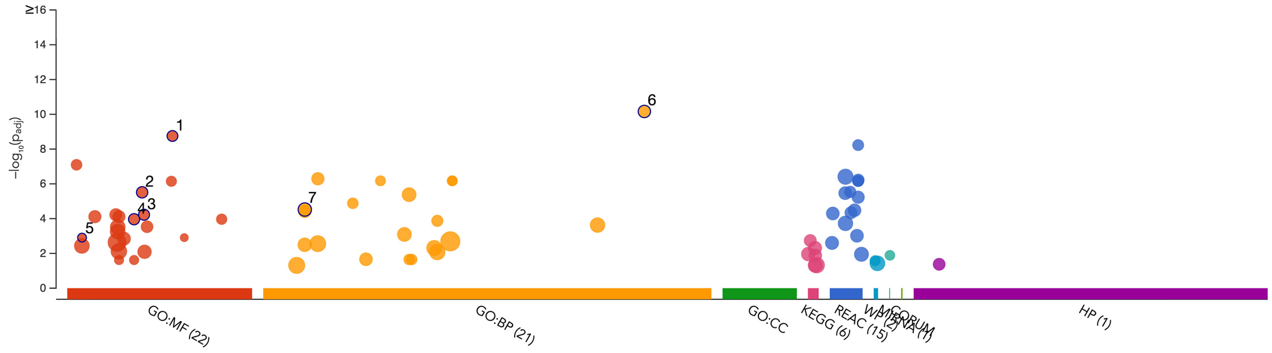


| ID | Source | Term ID | Term Name | $p_{adj}$ (query_1) |
|---|---|---|---|---|
| 1 | GO:MF | GO:0047086 | ketosteroid monooxygenase activity | $1.767 \times 10^{-9}$ |
| 2 | GO:MF | GO:0032052 | bile acid binding | $3.064 \times 10^{-6}$ |
| 3 | GO:MF | GO:0033130 | acetylcholine receptor binding | $5.980 \times 10^{-5}$ |
| 4 | GO:MF | GO:0030548 | acetylcholine receptor regulator activity | $1.083 \times 10^{-4}$ |
| 5 | GO:MF | GO:0004507 | steroid 11-beta-monooxygenase activity | $1.241 \times 10^{-3}$ |
| 6 | GO:BP | GO:1902644 | tertiary alcohol metabolic process | $6.879 \times 10^{-11}$ |
| 7 | GO:BP | GO:0006693 | prostaglandin metabolic process | $2.999 \times 10^{-5}$ |

**Supplementary Figure 2:** Manhattan plot of the g:GOSt GSORA outputs from the unsmoothed CSS analysis results (324 genes). Only significant functional terms are shown (g:SCS $P_{adj.} < 0.05$), and the functional entities are from the Gene Ontology (GO), KEGG, Reactome, WikiPathways (WP), miRTarBase, CORUM, and Human Phenotype Ontology (HP). The GO driver terms are numbered and shown with the g:SCC $P_{adj.}$ values for these GO terms (for further methodological details see: Kolberg *et al.* 2023).

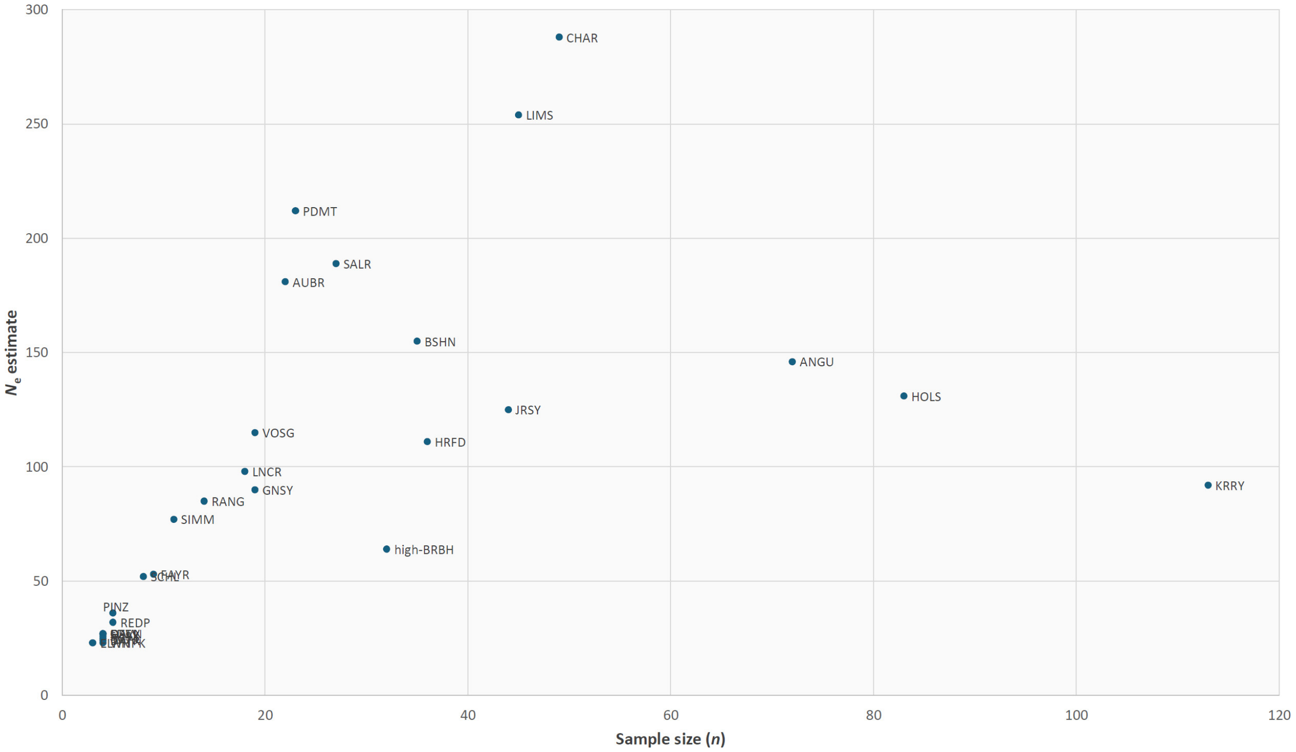


**Supplementary Figure 3:** Plot of effective population size ($N_e$) estimate versus sample size ($n$) for 29 European cattle breeds in the study.

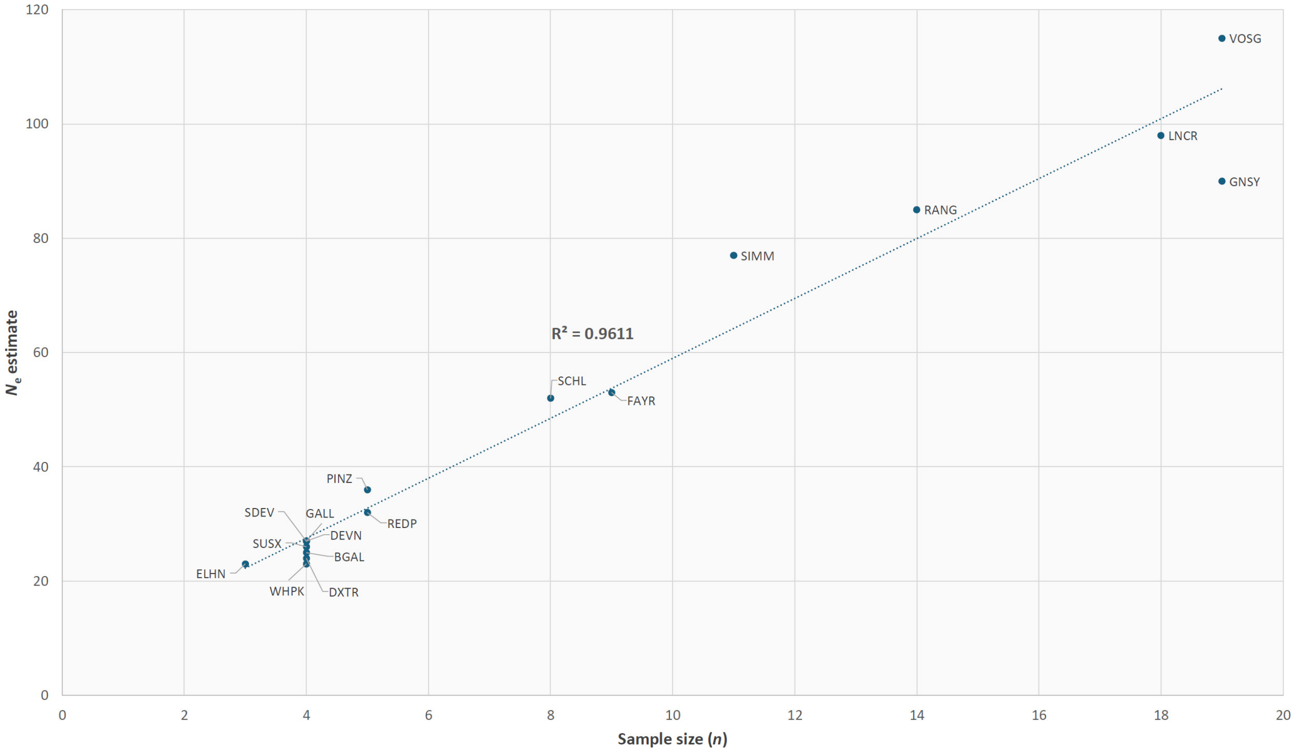


**Supplementary Figure 4:** Plot of effective population size ($N_e$) estimate versus sample size ($n$) for 17 European cattle breeds with $n$ < 20. A regression line is plotted with the coefficient of determination ($R^2$) indicated.

### Supplementary Tables

**Supplementary Table 1:** Breeds and populations used in this study, with sample size (*n*), purpose, country of origin, mean observed heterozygosity ($H_o$) and number of SNPs used for effective population size ($N_e$) modelling.

| Breed | Code | *n* | Purpose | Country of origin | Mean $H_o$ | No. of SNPs: $N_e$ modelling |
|---|---|---|---|---|---|---|
| Aberdeen Angus | ANGU | 72 | Beef | Scotland | 0.3380 | 19,637 |
| Aubrac | AUBR | 22 | Dual-purpose | France | 0.3447 | 22,298 |
| Belted Galloway | BGAL | 4 | Beef | Scotland | 0.3216 | 19,307 |
| BóRiabhach[a] | BRBH | 84 | Dual-purpose | Ireland | 0.3700 | 18,741 |
| British Shorthorn | BSHN | 35 | Dual-purpose | England | 0.2968 | 17,889 |
| Charolais | CHAR | 49 | Beef | France | 0.3622 | 20,761 |
| Devon | DEVN | 4 | Beef | England | 0.3170 | 19,127 |
| Dexter | DXTR | 4 | Dual-purpose | Ireland | 0.2722 | 18,361 |
| English Longhorn | ELHN | 3 | Dual-purpose/draught | England | 0.2452 | 13,467 |
| Finnish Ayrshire | FAYR | 9 | Dairy | Scotland/Finland | 0.3480 | 22,408 |
| Galloway | GALL | 4 | Beef | Scotland | 0.3257 | 19,115 |
| Guernsey | GNSY | 19 | Dairy | Channel Islands | 0.3140 | 21,577 |
| Hereford | HRFD | 36 | Beef | England | 0.3261 | 16,116 |
| Holstein | HOLS | 83 | Dairy | The Netherlands | 0.3671 | 22,304 |
| Jersey | JRSY | 44 | Dairy | Channel Islands | 0.3103 | 20,734 |
| Kerry | KRRY | 113 | Dairy | Ireland | 0.3387 | 9,234 |
| Lincoln Red | LNCR | 18 | Beef | England | 0.2919 | 18,166 |
| Limousin | LIMS | 45 | Beef/draught | France | 0.3536 | 23,123 |
| N'Dama | NDAG | 22 | Dual-purpose | Guinea (West Africa) | 0.2383 | --- |
| Piedmontese | PDMT | 23 | Dual-purpose | Italy | 0.3677 | 23,519 |
| Pinzgauer | PINZ | 5 | Dual-purpose | Austria | 0.3537 | 21,579 |

| | | | | | | |
|---|---|---|---|---|---|---|
| Red Angus | RANG | 14 | Beef | Scotland | 0.3445 | 22,158 |
| Red Poll | REDP | 5 | Beef | England | 0.3228 | 20,249 |
| Salers | SALR | 27 | Dual-purpose | France | 0.3274 | 21,387 |
| Scottish Highland | SCHL | 8 | Beef | Scotland | 0.3117 | 19,684 |
| Simmental | SIMM | 11 | Dual-purpose/draught | Switzerland | 0.3538 | 16,400 |
| South Devon | SDEV | 4 | Beef | England | 0.3362 | 21,574 |
| Sussex | SUSX | 4 | Beef/draft | England | 0.3092 | 18,633 |
| Vosgienne | VOSG | 19 | Dual-purpose | France | 0.3557 | 20,842 |
| White Park | WHPK | 4 | Dual-purpose/draught | England | 0.2502 | 15,698 |
| **TOTAL** | | **794** | | | | |

[a] $H_o$ is for all 84 BRBH animals, and the SNP number is for *high*-BRBH ($n$ = 28).

**Supplementary Table 2:** Overview of the number of SNPs and animals used for each analysis and the filters applied.

| **Analysis** | **Animals** | **Breeds/groups** | **SNPs** | **Filters and settings** |
|---|---|---|---|---|
| Merged BovineSNP50 dataset | 794 | 30 (29 European + NDAG) | 54,057 → 42,124 | Positions updated UMD3.1 → ARS-UCD1.2; autosomal SNPs with call rate < 95% removed |
| ADMIXTURE and PCA | 772 | 29 (NDAG excluded) | 13,653 | MAF ≥ 0.05 (25,399 SNPs), then LD pruning ($r^2$ < 0.1; 50-SNP window, 10-SNP step) |
| TreeMix | 738 | *high*-BRBH + 29 comparators (incl. NDAG) | 25,395 | MAF ≥ 0.05; SNPs genotyped in every population; blocks of 50 SNPs |
| SNeP ($N_e$), per breed | per breed | *high*-BRBH + 28 European breeds | per breed (**Supplementary Table 1**) | Within each breed, SNeP excluded SNPs with MAF < 0.05 or any missing genotype |
| ROH/$F_{ROH}$ | 764 | *high*-BRBH, low-BRBH + 28 European breeds | 25,346 | SNPs missing in > 50% of any breed removed; SNP call rate ≥ 95%; animals with > 10% missing genotypes removed; no MAF filter; no LD pruning |
| IBS and KING kinship | 84 | *high*-BRBH + low-BRBH | 13,875 | MAF ≥ 0.05; LD pruning ($r^2$ < 0.1; 50-SNP window, 10-SNP step) |
| CSS selection scan | 28 + 84 | *high*-BRBH versus HOLS | 24,380 | SNPs with values for all four component statistics ($\pi$, $F_{ST}$,Tajima's $D$, $\Delta SAF$) |

**Supplementary Table 3:** Ancestry components ($K$ = 1–9) for the 84 BóRiabhach (BRBH) cattle analysed in this study, with the BRBH ancestry component highlighted (***Supp_Table_3.xlsx***).

**Supplementary Table 4:** Modelled genetic effective population size ($N_e$) for 29 European production and traditional heritage cattle breeds (***Supp_Table_4.xlsx***).

**Supplementary Table 5:** Runs of homozygosity (ROH) detected in individual animals partitioned by length class. Also shown are the individual genomic inbreeding values ($F_{ROH}$) (***Supp_Table_5.xlsx***).

**Supplementary Table 6:** Genomic inbreeding (mean $F_{ROH}$) for individual breeds and population samples.

| **Breed** | ***n*** | **Mean $F_{ROH}$ (± SE)** |
|---|---|---|
| ANGU | 72 | 0.0646 ± 0.0042 |
| AUBR | 22 | 0.0192 ± 0.0043 |
| BGAL | 4 | 0.0797 ± 0.0100 |
| ***high*-BRBH** | 28 | 0.0393 ± 0.0121 |
| ***low*-BRBH** | 56 | 0.0144 ± 0.0025 |
| BSHN | 34 | 0.0949 ± 0.0058 |
| CHAR | 48 | 0.0219 ± 0.0024 |
| DEVN | 4 | 0.0694 ± 0.0198 |
| DXTR | 4 | 0.1868 ± 0.0532 |
| ELHN | 3 | 0.1801 ± 0.0125 |
| FAYR | 9 | 0.0413 ± 0.0048 |
| GALL | 4 | 0.0632 ± 0.0120 |
| GNSY | 19 | 0.0712 ± 0.0064 |
| HOLS | 83 | 0.0497 ± 0.0029 |
| HRFD | 35 | 0.1002 ± 0.0113 |

| **Breed** | ***n*** | **Mean $F_{ROH}$ (± SE)** |
|---|---|---|
| JRSY | 44 | 0.0658 ± 0.0042 |
| KRRY | 110 | 0.0612 ± 0.0023 |
| LIMS | 45 | 0.0190 ± 0.0028 |
| LNCR | 17 | 0.1035 ± 0.0090 |
| PDMT | 23 | 0.0053 ± 0.0012 |
| PINZ | 5 | 0.0233 ± 0.0064 |
| RANG | 14 | 0.0553 ± 0.0084 |
| REDP | 5 | 0.0878 ± 0.0102 |
| SALR | 27 | 0.0422 ± 0.0083 |
| SCHL | 8 | 0.0668 ± 0.0058 |
| SDEV | 3 | 0.0350 ± 0.0061 |
| SIMM | 11 | 0.0186 ± 0.0046 |
| SUSX | 4 | 0.0801 ± 0.0238 |
| VOSG | 19 | 0.0138 ± 0.0030 |
| WHPK | 4 | 0.2009 ± 0.0296 |

**Supplementary Table 7:** Pairwise genotype identity-by-state (IBS) matrix for all BRBH animals ($n$ = 84), with *high*-BRBH and *low*-BRBH animals highlighted in red and green text, respectively (individual animals are ordered by descending BRBH ancestry) (***Supp_Table_7.xlsx***).

**Supplementary Table 8:** Focal SNPs in unsmoothed selection signature peaks based on *CSS score* thresholding criteria (top 0.1% and at least one SNP in the top 1.0% within ±1 Mb).

| Chromosome | Genomic position | SNP ID | CSS score |
|---|---|---|---|
| 2 | 72,858,168 | 2:72858168 | 6.64 |
| 3 | 103,342,904 | 3:103342904 | 7.24 |
| 4 | 21,916,132 | 4:21916132 | 5.84 |
| 4 | 76,247,713 | 4:76247713 | 6.78 |
| 6 | 68,159,201 | 6:68159201 | 6.16 |
| 6 | 68,192,608 | 6:68192608 | 6.76 |
| 7 | 88,624,940 | 7:88624940 | 5.58 |
| 8 | 52,889,444 | 8:52889444 | 5.82 |
| 13 | 12,378,558 | 13:12378558 | 5.65 |
| 13 | 44,105,994 | 13:44105994 | 7.88 |
| 13 | 44,131,206 | 13:44131206 | 7.06 |
| 13 | 44,736,057 | 13:44736057 | 5.56 |
| 13 | 44,929,091 | 13:44929091 | 6.24 |
| 13 | 44,999,110 | 13:44999110 | 6.37 |
| 14 | 731,230 | 14:731230 | 6.04 |
| 16 | 43,266,548 | 16:43266548 | 5.55 |
| 16 | 43,291,228 | 16:43291228 | 5.55 |
| 16 | 49,659,723 | 16:49659723 | 5.56 |
| 16 | 59,841,996 | 16:59841996 | 6.87 |
| 20 | 39,383,171 | 20:39383171 | 6.09 |
| 20 | 42,113,227 | 20:42113227 | 8.41 |
| 21 | 40,888,569 | 21:40888569 | 6.62 |
| 22 | 2,329,624 | 22:2329624 | 5.57 |

**Supplementary Table 9:** Focal selection genomic windows identified from smoothed selection signature peaks based on *CSS score* thresholding criteria (top 0.1% within ±1 Mb).

| Chromosome | Genomic start position | Genomic end position | Number of SNPs | Maximum *CSS score* |
|---|---|---|---|---|
| 8 | 112,250,001 | 114,000,000 | 1 | 6.02 |
| 13 | 44,000,001 | 45,000,000 | 11 | 7.88 |
| 13 | 49,000,001 | 50,000,000 | 2 | 4.97 |
| 14 | 1 | 1,000,000 | 10 | 6.04 |
| 16 | 42,500,001 | 44,000,000 | 10 | 5.55 |

**Supplementary Table 10:** Genes contained within the selection peaks identified from the unsmoothed *CSS score* analysis (***Supp_Table_10.xlsx***).

**Supplementary Table 11:** Genes contained within the selection peaks identified from the smoothed *CSS score* analysis (***Supp_Table_11.xlsx***).

**Supplementary Table 12:** Functional terms identified from the GSORA performed with g:COSt in g:Profiler using the smoothed *CSS score* gene set (177 genes) (***Supp_Table_12.xlsx***).

**Supplementary Table 13:** Functional terms identified from the GSORA performed with g:COSt in g:Profiler using the unsmoothed *CSS score* gene set (324 genes) (***Supp_Table_13.xlsx***).
